# Emergence and features of the multipartite genome structure of the family *Burkholderiaceae* revealed through comparative and evolutionary genomics

**DOI:** 10.1101/382267

**Authors:** George C diCenzo, Alessio Mengoni, Elena Perrin

**Affiliations:** Department of Biology, University of Florence, Sesto Fiorentino, FI, 50019, Italy

**Keywords:** Bacterial genome evolution, Proteobacteria, divided genome structure, chromid, *Burkholderiaceae*, evolutionary biology, computational biology

## Abstract

The multipartite genome structure is found in a diverse group of important symbiotic and pathogenic bacteria; however, the advantage of this genome structure remains incompletely understood. Here, we perform comparative genomics of hundreds of finished β-proteobacterial genomes to study the role and emergence of multipartite genomes. Nearly all essential secondary replicons (chromids) of the β-proteobacteria are found in the family *Burkholderiaceae*. These replicons arose from just two plasmid acquisition events, and they were likely stabilized early in their evolution by the presence of core genes, at least some of which were likely acquired through an inter-replicon translocation event. On average, *Burkholderiaceae* genera with multipartite genomes had a larger total genome size, but smaller chromosome, than genera without secondary replicons. Pangenome-level functional enrichment analyses suggested that inter-replicon functional biases are partially driven by the enrichment of secondary replicons in the accessory pangenome fraction. Nevertheless, the small overlap in orthologous groups present in each replicon’s pangenome indicates a clear functional separation of the replicons. Chromids appeared biased to environmental adaptation, as the functional categories enriched on chromids were also over-represented on the chromosomes of the environmental genera (*Paraburkholderia, Cupriavidus*) compared to the pathogenic genera (*Burkholderia, Ralstonia*). Using ancestral state reconstruction, it was predicted that the rate of accumulation of modern-day genes by chromids was more rapid than the rate of gene accumulation by the chromosomes. Overall, the data are consistent with a model where the primary advantage of secondary replicons is in facilitating increased rates of gene acquisition through horizontal gene transfer, consequently resulting in a replicon enriched in genes associated with adaptation to novel environments.

## INTRODUCTION

The genomes of about 10% of bacterial species display a multipartite structure, consisting of a chromosome plus one or more additional replicons (Harrison et al. 2010; diCenzo and Finan 2017). The secondary replicons can be either essential (termed ‘chromid’) or non-essential (termed ‘megaplasmid’), and the primary features of these replicons and the multipartite genome structure has been recently reviewed (diCenzo and Finan 2017). Studies have indicated that the evolutionary trajectories (Galardini et al. 2013) as well as the rates of evolution (Cooper et al. 2010) and mutation (Dillon et al. 2015; Dillon et al. 2017) differ across replicons in a genome. Similarly, secondary replicons generally display higher variability at both the gene (Choudhary et al. 2007; Guo et al. 2009; Holden et al. 2009) and nucleotide (Chain et al. 2006; Bavishi et al. 2010; Epstein et al. 2012) level. Functional biases have also repeatedly been observed across replicons in a genome (Heidelberg et al. 2000; Chain et al. 2006; Janssen et al. 2010; diCenzo and Finan 2017), with chromosomes enriched in core functions and secondary replicons enriched in transport, metabolism, and regulatory functions. The reason for the evolution of the multipartite genome structure is not fully understood; however, its emergence may have been driven by adaptation to multiple niches (Chain et al. 2006; Galardini et al. 2013; diCenzo et al. 2014; diCenzo and Finan 2017).

Among the β-proteobacteria, multipartite genomes have been predominately identified in the genera *Burkholderia, Paraburkholderia, Cupriavidus*, and *Ralstonia* of the family *Burkholderiaceae* (Harrison et al. 2010; diCenzo and Finan 2017). These species generally possess large (5-10 Mb) genomes, all of them consisting of at least a chromosome and a chromid (Coenye 2014). The latter is often referred to as a second chromosome, and it can be nearly as large as the primary chromosome. Species belonging to the *Burkholderia cepacia* complex (Bcc) contain a third large replicon (~ 1 Mb) that for many years has been defined as a third chromosome. However, recent work has demonstrated that, at least in some Bcc species, this replicon is a non-essential megaplasmid involved in virulence and stress tolerance (Agnoli et al. 2012; Agnoli et al. 2014). Nevertheless, genome-scale mutageneses have suggested that all replicons in *Burkholderia cenocepacia* strains J2315 and H111 carry genes required for optimal fitness in laboratory conditions (Wong et al. 2016; Higgins et al. 2017). As inter-replicon (essential) gene transfer can occur in multipartite genomes (Guo et al. 2010; diCenzo et al. 2013), it is possible that the essentiality of the third replicon could be strain specific, as suggested by Wong et al. (Wong et al. 2016).

The family *Burkholderiaceae* represents an ideal model for studying the evolution and function of bacterial multipartite genomes. The species of the *Burkholderiaceae* contain numerous human and animal pathogens and opportunistic pathogens, including the genera *Burkholderia* and *Pandoraea* found in the lungs of cystic fibrosis patients (Schaffer 2015). This family also contains numerous plant pathogens (e.g., the genus *Ralstonia*) (Denny 2007), plant symbionts (Gyaneshwar et al. 2011), insect symbionts (Takeshita and Kikuchi 2017), and can be found in soil, water, and polluted environments (Coenye and Vandamme 2003). This high environmental plasticity, coupled with the public availability of hundreds of finished genomes, facilitates comparative genomic analyses and the identification of genes (and replicons) associated with these environments. Additionally, the family *Burkholderiaceae* contain species with and without multipartite genomes, enabling the identification of which features, if any, are associated with the presence of a multipartite genome.

In this work, we attempt to address open questions on bacterial multipartite genomes by employing a computational approach to characterize the distribution, emergence, and putative functions of multipartite genomes in the β-proteobacteria, focusing on 293 finished *Burkholderiaceae* genomes. Our results suggest that all chromids in the family *Burkholderiaceae* can be explained by two plasmid acquisition events, and that early gain of genes from the chromosome may have helped stabilize these replicons. Additionally, based on comparative genomics and ancestral state reconstruction analyses, we propose that the primary advantage provided by a secondary replicon is increased genetic malleability, which in turn results in the replicon accumulating niche specialized functions.

## MATERIALS AND METHODS

### Classification of bacterial replicons

The 960 genomes of the β-proteobacteria available through the National Center for Biotechnology Information (NCBI) Genome database (12/12/2017) that fit our criteria were downloaded (Data Set S1). The criteria for inclusion were as follows: i) assembly level of ‘complete’ or ‘chromosome’; ii) contained no unplaced scaffolds; and iii) had a RefSeq accession. Additionally, *Burkholderia* sp. PAMC 26561 was excluded as two replicons (NC_CP014308.1 and NC_CP014309.1) were almost identical, suggesting there was an error in the assembly. Replicons were classified as done elsewhere (diCenzo and Finan 2017). Briefly, the largest replicon in each genome was designated the chromosome, while replicons less than 350 kb were defined as a plasmid. Of the remaining replicons, those with a %GC content within 1% of the chromosome and with a dinucleotide relative abundance distance [calculated as described by (diCenzo and Finan 2017)] from the chromosome of less than 0.4 were annotated as a putative chromid, with all remaining replicons designated as a megaplasmid. The classifications and metadata for each replicon is available in Data Set S1, and the species names used are those given in NCBI.

A representative set of β-proteobacterial genomes was chosen based on the following criteria: i) assembly level of ‘complete’ or ‘chromosome’; ii) contained no unassigned scaffolds; iii) had a RefSeq accession; iv) had a genus and species name; and v) if there were multiple genomes available for a species, one representative was randomly chosen. This resulted in the identification of a set of 176 representative strains (Data Set S2).

A set of genomes from the family *Burkholderiaceae* was also prepared. The criteria for inclusion in this strain set was: i) assembly level of ‘complete’ or ‘chromosome’; ii) contained no unassigned scaffolds; iii) had a RefSeq accession; and iv) had a genus. The final list consisted of 293 *Burkholderiaceae* strains (Data Set S3). Within this family, the classification of bacterial replicons was refined based on known data; the largest non-chromosomal replicon in the genera *Burkholderia, Paraburkholderia, Ralstonia*, and *Cupriavidus* was termed a chromid, while other large replicons were termed megaplasmids (Chain et al. 2006; Agnoli et al. 2012).

### Phylogenetic reconstructions of the β-proteobacteria

A phylogeny of the β-proteobacteria was prepared based on the proteomes of the 176 representative strains. The phylogeny was prepared with our in-house pipeline (diCenzo et al. 2017). In brief, AMPHORA2 (Wu and Scott 2012) was used to identify the presence of 31 highly conserved proteins in each proteome using hidden Markov models (HMMs) and HMMER version 3.1b2 (Eddy 2009). The 26 proteins (Frr, InfC, NusA, RplB, RplC, RplD, RplE, RplF, RplK, RplL, RplM, RplN, RplP, RplS, RplT, RpmA, RpoB, RpsB, RpsC, RpsE, RpsI, RpsJ, RpsK, RpsM, RpsS, Tsf) found in at least 95% of the genomes and present in single-copy in each of those genomes were collected. Each set of orthologs was aligned using Mafft version 7.310 (Katoh and Standley 2013) with the ‘localpair’ option and with 40 threads. Alignments were trimmed using TrimAl version 1.2rev59 (Capella-Gutiérrez et al. 2009) and the ‘automated1’ option, and the alignments were then concatenated. The RAxML version 8.2.10 algorithm (Stamatakis 2014) on the Cipres webserver (Miller et al. 2010) was used to build a maximum likelihood phylogeny, using the LG amino acid substitution model and the CAT rate heterogeneity. The final tree is the bootstrap best tree following 108 bootstrap replicates, which was visualized using FigTree (http://tree.bio.ed.ac.uk/software/figtree/).

A phylogeny of the family *Burkholderiaceae* was constructed based on the proteomes of all 293 *Burkholderiaceae* strains. The procedure was essentially the same as described above for the 176 representative β-proteobacteria strains. The only differences were: i) the alignment was based on 28 proteins (Frr, InfC, NusA, RplA, RplB, RplC, RplD, RplE, RplF, RplK, RplL, RplM, RplN, RplP, RplS, RplT, RpmA, RpoB, RpsB, RpsC, RpsE, RpsI, RpsJ, RpsK, RpsM, RpsS, SmpB, Tsf); and ii) 360 bootstrap replicates were performed.

A third phylogeny was constructed based on the proteomes of all 973 β-proteobacteria strains (Figure S1). The procedure was performed largely as described above for the 176 representative β-proteobacteria strains. The differences were: i) the alignment was based on only 24 proteins (Frr, InfC, NusA, RplC, RplD, RplE, RplK, RplL, RplM, RplN, RplP, RplS, RplT, RpmA, RpoB, RpsB, RpsC, RpsE, RpsI, RpsJ, RpsK, RpsM, RpsS, Tsf); and ii) 1008 bootstrap replicates were performed.

### Replicon phylogenetic analyses

Phylogenies of all replicons of the family *Burkholderiaceae* were built based on the partitioning protein (ParB) and the replication protein (Rep). The seed alignment files of the ParBc and of the Rep_3 protein families were downloaded from Pfam (Finn et al. 2016), and HMMs for each were built with the hmmbuild function of HMMER. Additionally, the complete Pfam-A version 31.0 (16,712 HMMs) and TIGERFAM version 15.0 (4,488 HMMs) databases were downloaded (Haft et al. 2013; Finn et al. 2016). The hmmconvert function of HMMER was run on both databases to ensure consistent formatting, following which the two databases were concatenated and hmmpress of HMMER used to prepare a hmmscan searchable database.

All proteins of all replicons from 285 of the 293 strains of the *Burkholderiaceae* were combined as a single fasta file, and the proteins were renamed to keep track of which replicon they originated from. The eight strains in which multiple replicons were integrated as a single replicon in the strain’s genome assembly were excluded from the analysis. Both HMMs (ParBc and Rep_3) were individually searched against the all-strains proteome using the hmmsearch function of HMMER. A custom Perl script was used to parse the hmmsearch output files from both searches, and to prepare a fasta file with all hits regardless of e-value (one file for ParBc and one for Rep_3). The fasta files were then individually searched against the combined HMM database described above with the hmmscan function of HMMER. A Perl script was prepared and used to parse the hmmscan output and identify the top scoring HMM hit for each protein. Proteins were annotated as ParB if the top hit was ParBc (Pfam), RepB (Pfam), or TIGR00180 (TIGRFAM). Proteins were annotated as Rep if the top hit was either Rep_3 (Pfam) or RPA (Pfam). Finally, all orthologs were combined as a single fasta file.

For each of ParB and Rep, the proteins were aligned using Mafft with the ‘localpair’ option and with 40 threads, and the alignments then trimmed using TrimAl with the ‘automated1’ option. Maximum likelihood phylogenies were built using the RAxML algorithm on the Cipres webserver, with the LG amino acid substitution model and the CAT rate heterogeneity. The final trees are the bootstrap best trees following 504 or 756 bootstrap replicates for ParB or Rep, respectively.

### Synteny analysis

Replicon nucleotide fasta files were downloaded from NCBI. Pairwise comparisons of replicons were performed with nucmer version 3.1 of the mummer package (Kurtz et al. 2004), using default parameters and the ‘-p’ option. Alignments from nucmer where used for construction of dot plots using the mummerplot version 3.5 function of the mummer package, using default parameters and the ‘-f’ and ‘-l’ options.

### Average nucleotide identity (ANI) calculations

All-against-all pairwise ANI values for all 293 genomes of the family *Burkholderiaceae* were calculated using FastANI with default parameters (Jain et al. 2017). The output of FastANI was used to construct an ANI matrix (Figure S2), and in cases where no value was returned by FastANI (due the percent identity being too low), a value of zero was added. The matrix was clustered and visualized in R using the heatmap.2 function of the gplots package (Warnes et al. 2009), with clustering performed using average linkage.

### Pangenome calculation and core genome comparison

Prior to pangenome analyses, all genomes were re-annotated using Prokka version 1.12-beta (Seemann 2014) with the ‘fast’, ‘norrna’, and ‘notrna’ options. This was done to ensure consistent annotation and to prepare a GFF file with the correct format for use with Roary. A custom Perl script was used to prepare replicon-specific GFF files from the GFF and FNA files returned by Prokka. All pangenomes were calculated using Roary version 3.7.0 (Page et al. 2015) with 40 threads, the ‘-e’, ‘-n’, and ‘-g 100000’ options, and with an identity threshold of 80%. Core genes were identified as those found in at least 99% of the genomes/replicons.

In calculating the pangenomes, the following genomes were included in the genus *Paraburkholderia:* i) all genomes annotated with *Paraburkholderia* as the genus, except for *Paraburkholderia rhizoxinica* HKI 454; and ii) the *Burkholderia* spp. CCGE1101, CCGE1102, CCGE1103, HB1, OLGA172, PAMC 26561, PAMC 28687, RPE64, RPE67, KK1, and Y123, as these grouped with the *Paraburkholderia* in both the phylogeny and ANI matrix. *P. rhizoxinica* HKI 454 was excluded on the basis of the phylogeny and previous work (Sawana et al. 2014), which suggest this strain may belong to its own genus separate from both the *Burkholderia* and the *Paraburkholderia*. All strains annotated with the genus *Burkholderia* not included in the genus *Paraburkholderia* were included in the calculation of the pangenome of the genus *Burkholderia*. In the case of *Cupriavidus*, the following genomes were included: i) all genomes annotated with *Cupriavidus* as the genus, and ii) *Ralstonia pickettii* DTP0602, *Ralstonia eutropha* H16, and *R. eutropha* JMP134, as these grouped with the *Cupriavidus* in both the phylogeny and ANI matrix. All strains annotated with the genus *Ralstonia* not included in the genus *Cupriavidus* were included in the calculation of the pangenome of the genus *Ralstonia*.

To compare the core genomes of the different genera, a Blast bidirectional best hit (Blast-BBH) approach was employed. For each pair-wise comparison, the nucleotide pan_genome_reference.fa files returned by Roary were converted into amino acid fasta files using transeq of the EMBOSS version 6.6.0.0 package (Rice et al. 2000). The resulting proteomes were searched against each other using Blastp of the BLAST 2.2.29+ package (Camacho et al. 2009) with the options: culling_limit 1, max_target_seqs 1, max_hsps 1, and 32 threads. A custom shell pipeline was used to identify putative orthologs as proteins whose top hit was each other in the Blastp searches. A custom shell pipeline was then used to identify which pairs of orthologs belonged to the core gene set of both genera.

### McDonald-Kreitman test

McDonald-Kreitman tests (McDonald and Kreitman 1991) were performed using an online webserver (http://mkt.uab.es/mkt/; (Egea et al. 2008)). The ParB proteins from each of the chromids for all *Burkholderia* species outside of the Bcc were used as ‘species 1’, while the ParB proteins from each of the chromids for all Bcc species were used as ‘species 2’.

### Functional analyses

Functional annotation of the accessory pangenomes was performed on the replicon-specific pangenomes produced as described above. A custom pipeline using Bash, Perl, and MATLAB (mathworks.com) scripts was used to collect the representative genes for each orthologous protein family present in less than 95% of strains, based on the ‘gene_presence_absence.csv’ and ‘pan_genome_reference.fa’ files produced by Roary (Page et al. 2015). Each coding sequence was converted to the corresponding amino acid sequence using the transeq function of the EMBOSS package (Rice et al. 2000). Eggnog-mapper (Huerta-Cepas et al. 2017) was then used to functionally annotate the proteins using the ‘bact’ database, 32 CPUs, and the ‘usemem’ option. Next, a Bash script was used to extract the eggnog-mapper output for just those proteins present in less than 75% or 25% of the strains, as well as to count the total number of proteins present in less than 75% or 25% of the strains. The collected data was then parsed using a Perl script, which returned the number or proteins and percentage of proteins annotated with each COG category (Tatusov et al. 2003). Fisher exact tests for pairwise comparisons of the abundance of each COG category in different pangenome sets were calculated in MATLAB. Chi-squared tests were performed manually in Excel. The number of eggNOG orthologous groups represented within each COG category was determined using a custom MATLAB script.

### Ancestral state reconstruction

All proteins from the 293 *Burkholderiaceae* strains were grouped into orthologous clusters using cd-hit version 4.6 (Li and Godzik 2006), with an identity cut-off of 40% (-c 0.4), with an alignment length of at least 70% of the shorter protein (-aS 0.7), and following additional options: −G 0, −T 30, −n 2, −d 0, and −M 50000. A total of 63,554 clusters were produced. The output of cd-hit was converted into a binary presence (one) and absence (zero) matrix, indicating which clusters were found in which strains.

Each of the 63,554 clusters were associated to the Roary produced pangenomes, described above, for the chromosomes and chromids of each of the genera *Burkholderia, Paraburkholderia, Ralstonia*, and *Cupriavidus*. The strains were then split into two groups: i) the *Burkholderia* and *Paraburkholderia* strains, and ii) the *Ralstonia* and *Cupriavidus* strains. In both groups, the chromosomal clusters of the two genera were merged as a single list, as were the chromid clusters. Then, clusters found on the chromosomes but not chromids of that group were identified, and the clusters specific to the chromids and absent on the chromosomes of that group were also identified. For the *Burkholderia/Paraburkholderia* group, there were 13,625 chromosome clusters and 12,332 chromid clusters. For the *Cupriavidus/Ralstonia* group, there were 9,908 chromosome clusters and 6,474 chromid clusters.

Ancestral state reconstruction was performed using the *ace* function of the APE package version 5.1 implemented in R (Paradis et al. 2004). This was done using the matrix produced based on the cd-hit output, and the phylogeny of the 293 *Burkholderiaceae* strains rooted based on the β-proteobacteria phylogeny of Figure 1. Ancestral state reconstruction was run independently for each cluster, using the equal rates model. If the probability of the ancestor having the gene was above 50%, the strain was said to contain the cluster; otherwise, the cluster was said to be absent in that strain. Next, for each set of clusters of interest (e.g., *Burkholderia/Paraburkholderia* chromosomal clusters), the number of clusters predicted to be present in the ancestor were summed. The summed ancestral reconstruction data were then mapped to the phylogeny using the phytools package version 0.6.44 of R (Revell 2011).

**Figure 1.**
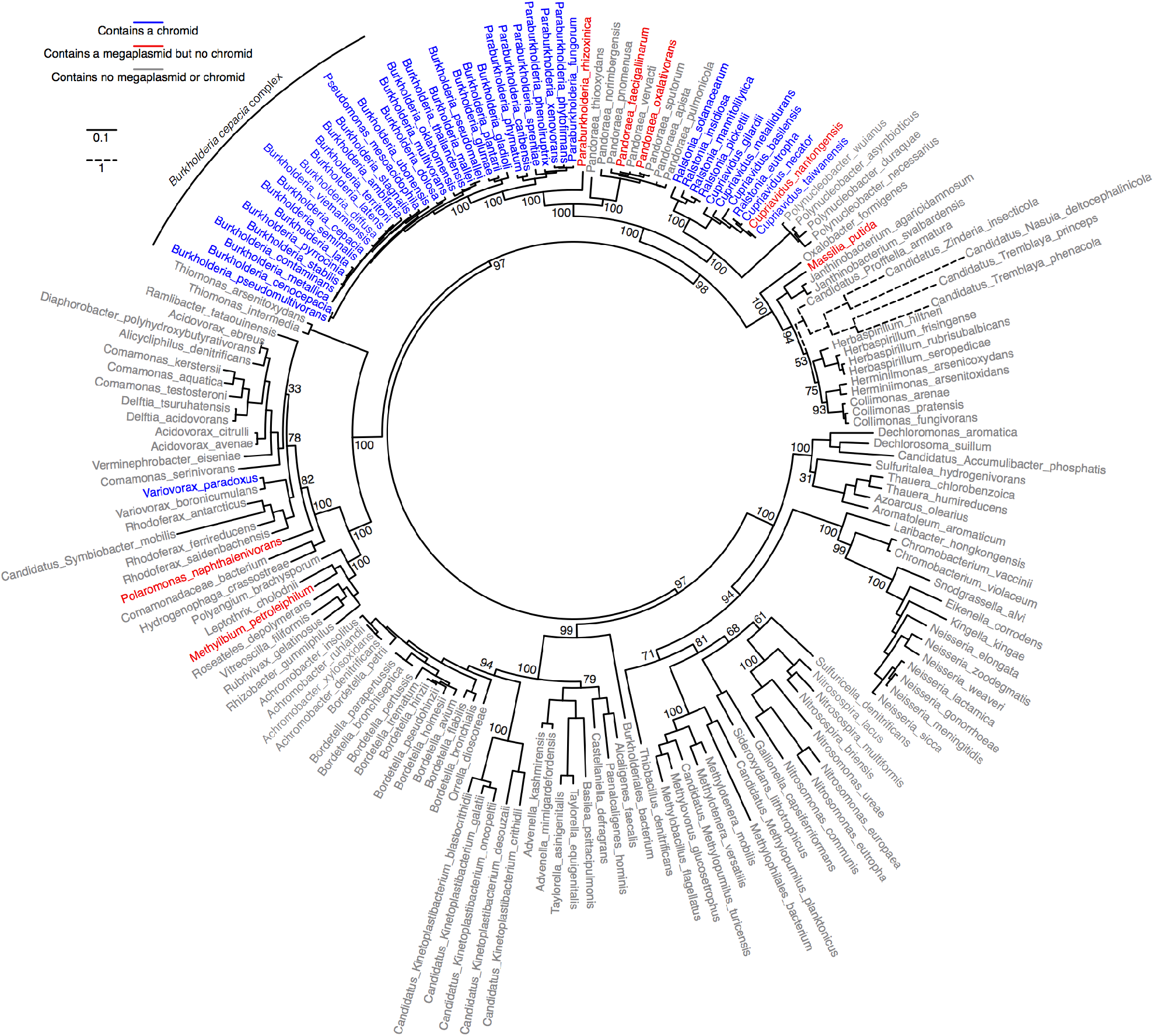
Distribution of multipartite genomes in the β-proteobacteria. A RAxML maximum likelihood phylogeny of the β-proteobacteria based on 26 highly conserved proteins (see methods). A colour legend is provided in the figure. Species with a putative chromid (blue) may also contain a megaplasmid. The branch lengths for one clade were reduced for presentation purposes; this clade is in dashed lines, and a separate scale is provided. Bootstrap values based on 108 replicates are provided where space permits. The original figure, the Newick formatted phylogeny, and the annotation file are provided in Data Set S6.

### Classification of species’ lifestyles

Information on the lifestyle of species of the family *Burkholderiaceae* was recovered from the GOLD database (Mukherjee et al. 2017), from the Global Catalogue of Microorganisms (Wu et al. 2013) and from literature source. A species was said to exhibit a certain lifestyle when at least one strain of that species was isolated from a given environment.

## RESULTS AND DISCUSSION

### Distribution of multipartite genomes in the β-proteobacteria

We began by examining the genome organization of representatives of 176 named species of the β-proteobacteria with a complete genome (see Materials and Methods). Genomes were considered multipartite if they contained two or more replicons of at least 350 kb in size; secondary replicons were classified into chromids and megaplasmids on the basis of GC content and dinucleotide composition (Data Set S2). Nearly all of the multipartite genomes found in this class of bacteria are within two clades of the family *Burkholderiaceae* (Figure 1). One of these clades consists of the genera *Burkholderia* and *Paraburkholderia*, the other includes the genera *Cupriavidus* and *Ralstonia*. Outside of these four genera, megaplasmids are found in two of the nine *Pandoraea* species (family *Burkholderiaceae*), while only three of the other 127 β-proteobacterial species had a megaplasmid (*Methylibium petroleiphilum, Polaromonas naphthalenivorans*, *Massilia putida*). Additionally, a chromid is found in only one species (*Variovorax paradoxus*) outside of the family *Burkholderiaceae*; the presence of a chromid in *Variovorax* sp. WDL1 has also been recently reported (Albers et al. 2018).

### Genome organization in the family *Burkholderiaceae*

As the large majority of the β-proteobacterial species with a multipartite genome are within the family *Burkholderiaceae*, we chose to focus our analyses on 293 genomes from this family (see Methods; Data Set S3). Examining the size distribution of the secondary replicons of this family revealed four main size groupings (Figure S3). Notably, replicons larger than 800 kb tend to be widespread within a clade of species, whereas those smaller than 800 kb tend to be strain-or species-specific. Therefore, we further focused our analyses of secondary replicons to those greater than 800 kb in size, expanded to include smaller replicons displaying clear signs of synteny to a secondary replicon larger than 800 kb. These large replicons were classified as large megaplasmids and chromids based on the published literature (Salanoubat et al. 2002; Chain et al. 2006; Fricke et al. 2009; Harrison et al. 2010; Agnoli et al. 2012; Higgins et al. 2017).

Three primary genome architectures are found within the family *Burkholderiaceae* (Figure 2 and Table 1). Species belonging to the genera *Polynucleobacter* and *Pandoraea* contain genomes largely present as a single chromosome, lacking large secondary replicons. In contrast, all species of the sister genera *Cupriavidus* and *Ralstonia* contain a chromid and lack large megaplasmids, with the exception *Cupriavidus necator* N-1, which has a large megaplasmid. Similarly, nearly all species of the sister genera *Burkholderia* and *Paraburkholderia* contain a chromid. The exception is *Paraburkholderia rhizoxinica* HKI 454; this strain, which contains a large megaplasmid, is an outgroup of the *Burkholderia* and *Paraburkholderia* clade, consistent with recent work suggesting that *P. rhizoxinicia* should be moved to its own genus (Estrada-de los Santos et al. 2013; Beukes et al. 2017). Many of the *Burkholderia* and *Paraburkholderia* species additionally contain a large megaplasmid. In the *Burkholderia*, the presence of a large megaplasmid is specific to the *Burkholderia cepacia* complex (Bcc; see Figure 1 for an indication of the species belong to the BCC). In the case of the genus *Paraburkholderia*, approximately half of the genomes contain a large megaplasmid, but these strains do not form a monophyletic group. Moreover, a monophyletic group of four strains, corresponding to the stink-bug-associated beneficial and environmental group (Takeshita and Kikuchi 2017), contain a second large megaplasmid; strikingly, the chromosome accounts for only 33-43% of the total genome size of these strains (Table 1).

**Figure 2.**
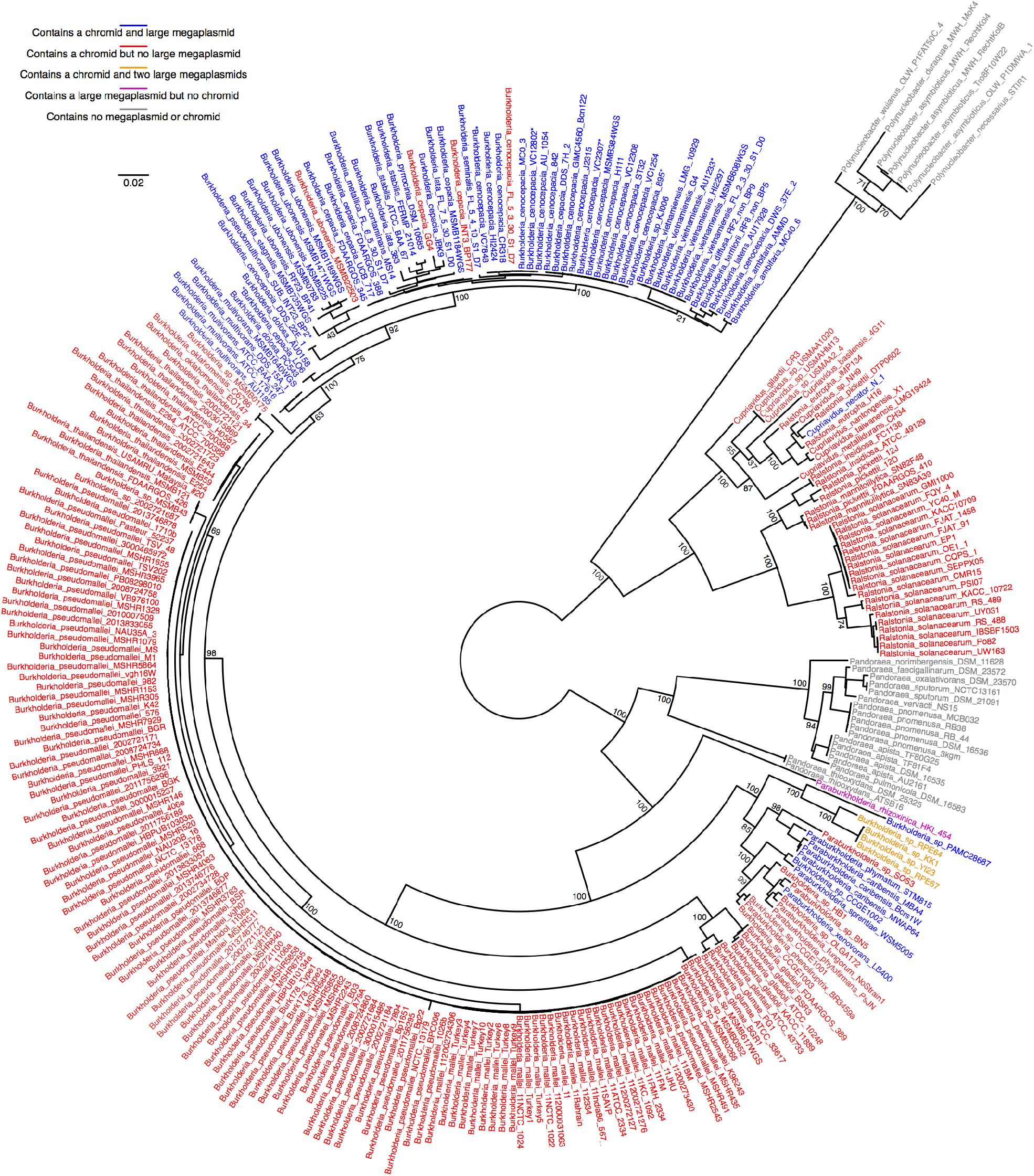
Genome organization within the family *Burkholderiaceae*. A RAxML maximum likelihood phylogeny of the β-proteobacteria based on 28 highly conserved proteins (see methods). A colour legend is provided in the figure. Strains marked with an asterisks (*) contain genomic regions homologous the chromosome, chromid, and megaplasmid, but with all or part of at least two of the replicons present as a co-integrant based on the assembly available through NCBI. Bootstrap values based on 360 replicates are provided where space permits. The original figure, the Newick formatted phylogeny, and the annotation file are provided in Data Set S6.

**Table 1.** Median genome and replicon sizes in the family *Burkholderiaceae*. Values indicate the median ± standard error based on the number of strains indicated in the second column. The section ‘All strains’ is based on all genome sequences within the clade, whereas the ‘Representative strains’ is based on one representative for each species within the clade. The megaplasmid, and 2^nd^ megaplasmid, only refer to the large (> 800 kb) megaplasmids, and does not consider additional, smaller replicons. The total genome size is based on all replicons in the genome.

| Genus or clade | Strain Count | Total | Chromosome | Chromid | Megaplasmid | 2<sup>nd</sup> Megaplasmid |
| --- | --- | --- | --- | --- | --- | --- |
| All strains |  |  |  |  |  |  |
| *Burkholderia cepacia complex* | 54 | 7.66 ± 0.10 | 3.55 ± 0.02 | 3.03 ± 0.06 | 1.05 ± 0.03 | - |
| Other *Burkholderia* species | 147 | 7.23 ± 0.06 | 4.02 ± 0.02 | 3.15 ± 0.04 | - | - |
| *Paraburkholderia* with 2 megaplasms | 4 | 8.79 ± 0.58 | 3.11 ± 0.10 | 1.78 ± 0.10 | 1.63 ± 0.19 | 1.50 ± 0.14 |
| *Paraburkholderia* with megaplasmid | 8 | 9.18 ± 0.58 | 3.61 ± 0.17 | 2.79 ± 0.21 | 1.80 ± 0.20 | - |
| *Paraburkholderia* without megaplasms | 9 | 7.65 ± 0.26 | 4.38 ± 0.07 | 2.99 ± 0.11 | - | - |
| *Cupriavidus* * | 13 | 7.82 ± 0.25 | 4.18 ± 0.11 | 2.89 ± 0.15 | - | - |
| *Ralstonia* | 26 | 5.61 ± 0.07 | 3.62 ± 0.04 | 1.98 ± 0.06 | - | - |
| *Pandoraea* | 18 | 5.60 ± 0.12 | 5.58 ± 0.10 | - | - | - |
| *Polynucleobacter* | 7 | 2.13 ± 0.11 | 2.13 ± 0.11 | - | - | - |
| Representative strains |  |  |  |  |  |  |
| *Burkholderia cepacia complex* | 17 | 7.58 ± 0.17 | 3.56 ± 0.04 | 3.00 ± 0.10 | 1.09 ± 0.05 | - |
| Other *Burkholderia* species | 8 | 7.30 ± 0.33 | 4.08 ± 0.16 | 3.06 ± 0.18 | - | - |
| *Paraburkholderia* with 2 megaplasms | 0 | - | - | - | - | - |
| *Paraburkholderia* with megaplasmid | 4 | 8.85 ± 0.39 | 3.67 ± 0.32 | 2.79 ± 0.16 | 1.69 ± 0.23 | - |
| *Paraburkholderia* without megaplasms | 3 | 8.21 ± 0.41 | 4.46 ± 0.10 | 3.51 ± 0.29 | - | - |
| *Cupriavidus* * | 6 | 7.14 ± 0.39 | 3.93 ± 0.17 | 2.58 ± 0.22 | - | - |
| *Ralstonia* | 4 | 5.33 ± 0.26 | 3.55 ± 0.13 | 1.72 ± 0.16 | - | - |
| *Pandoraea* | 9 | 5.74 ± 0.19 | 5.63 ± 0.16 | - | - | - |
| *Polynucleobacter* | 4 | 2.13 ± 0.17 | 2.13 ± 0.17 | - | - | - |

\* *Cupriavidus necator* N-1 has a 1.5 Mb megaplasmid.

### Phylogenetic analysis of the *Burkholderiaceae* secondary replicons

Phylogenies of all replicons from the family *Burkholderiaceae* were prepared as a means to evaluate the evolutionary relationship between the secondary replicons of this family. Phylogenies were prepared on the basis of the amino acid sequences of the partitioning protein ParB (Figure 3) and of the replication protein Rep (Figure S4). As described in the Methods, ParB proteins were identified based on similarity to the ParBc hidden Markov model (HMM) of Pfam, while Rep proteins were identified based on the Rep_3 HMM of Pfam (Finn et al. 2016). All phylogenies are consistent with those reported previously (Fricke et al. 2009; Lykidis et al. 2010; Passot et al. 2012). Synteny and pangenome analyses of the replicons were also performed to complement the phylogenetic data.

**Figure 3.**
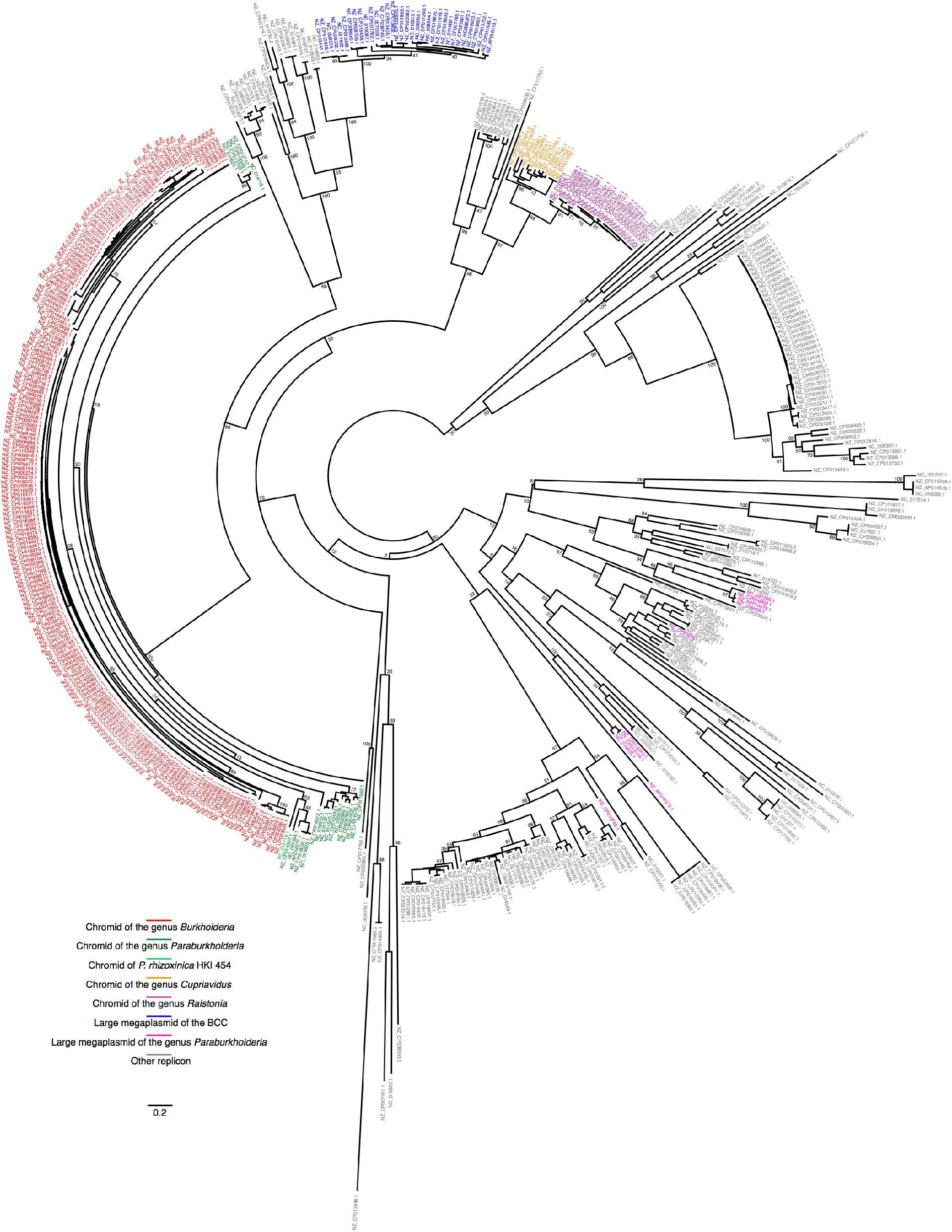
Phylogenetic relationship of the replicons of the family *Burkholderiaceae*. A RAxML maximum likelihood phylogeny of the replicons of the family *Burkholderiaceae* based on the amino acid sequence of the ParB partitioning protein. Each taxon is named according to the NCBI accession number of the replicon. A colour legend is provided in the figure. Bootstrap values based on 504 replicates are provided where space permits. The original figure, the Newick formatted phylogeny, and the annotation file are provided in Data Set S6.

### *The* Burkholderia *and* Paraburkholderia *chromids share common ancestry*

The chromids of the genera *Burkholderia* and *Paraburkholderia* form a clear cluster in both the ParB and Rep phylogenies (Figures 3 and S4). The chromids of the genus *Burkholderia* form a monophyletic group separate from those of the genus *Paraburkholderia* in the ParB phylogeny, whereas the chromids of the genus *Paraburkholderia* were nested within the chromids of the genus *Burkholderia* in the Rep phylogeny. The ParB and Rep proteins of the megaplasmid of *P. rhizoxinica* HKI 454, which appears to lack a chromid (Figure 2), also groups with chromids of the genera *Burkholderia* and *Paraburkholderia*. Additionally, a BLASTp search of the NCBI non-redundant protein database with the ParB protein of *Burkholderia lata* 383 chromid identified closely related proteins in species of the genus *Caballeronia* (> 98% coverage, and > 75% identity). The genus groups together with the genera *Burkholderia* and *Paraburkholderia* (Dobritsa and Samadpour 2016; Beukes et al. 2017). These results suggest that all chromids of the *Burkholderia/Paraburkholderia/Caballeronia* clade are derived from a single replicon acquired by an ancestor at the base of this taxon prior to its divergence.

The above conclusion is supported by synteny and gene content analyses. Although there is significant genetic variation, clear stretches of synteny can be seen when comparing between chromids of the genus *Burkholderia* as well as when comparing chromids of the genus *Paraburkholderia* (Figure S5). Additionally, a core set of chromid genes (on at least 99% of the chromids) is observed for both genera: 130 genes belong to the core gene set of the genus *Burkholderia*, while 74 genes are part of the core gene set of the genus *Paraburkholderia* (Table 2, Data Set S5). Moreover, 37 of the core genes are common between the *Burkholderia* and the *Paraburkholderia* (Table 2, Data Set S5). These results thus support that the chromids of all species within these genera share common ancestry. However, the low level of synteny and gene conservation may suggest that the ancestral plasmid contained relatively few genes found on the modern day chromids.

**Table 2.** Replicon-specific pangenome summary statistics.

| Genus or group | Replicon Type | Total Genes | Core Genes |
| --- | --- | --- | --- |
| *Burkholderia* | Chromid | 32,023 | 130 |
| *Paraburkholderia* | Chromid | 21,308 | 74 |
| *Burkholderia-Paraburkholderia* | Chromid | ND * | 37 |
| *Cupriavidus* | Chromid | 16,082 | 147 |
| *Ralstonia* | Chromid | 6,976 | 304 |
| *Cupriavidus-Ralstonia* | Chromid | ND | 58 |
| Bcc | Megaplasmid | 10,004 | 18 |

\*ND: Not determined

The 37 genes common to the chromids of *Burkholderia* and *Paraburkholderia* include genes involved in fatty acid and folate biosynthesis, purine, propanoate and amino acid metabolism, oxidative phosphorylation, and the citrate cycle (TCA cycle). In addition, genes coding for a DNA primase, a DNA polymerase I, and the RNA polymerase sigma factor RpoD are also present on the chromid. Approximately half of the 37 conserved genes belong to a large gene cluster that has been suggested to have moved from the chromosome to the chromid in an ancestral strain (Higgins et al. 2017). Additionally, eight of the conserved genes (*accD, asd, dnaG, dxs, folC, rpoD, sdhA* and *sdhD*) that belong to this gene cluster have been annotated as essential in at least one *Burkholderia* species based on genome-wide screens (Baugh et al. 2013; Moule et al. 2014; Wong et al. 2016; Higgins et al. 2017). The presence of this essential gene region on nearly all examined *Burkholderia* and *Paraburkholderia* chromids suggests that the ancestral plasmid giving rise to the modern-day chromids acquired this region early after its capture by the ancestor of the *Burkholderia/Paraburkholderia*. Thus, this inter-replicon translocation of an essential gene region likely helped stabilize the ancestral plasmid, contributing to its maintenance and emergence as a chromid.

### *The* Cupriavidus *and* Ralstonia *chromids share common ancestry*

A similar situation as above was observed for the genera *Ralstonia* and *Cupriavidus*. The chromids of all species from these genera form a single clade in the ParB phylogeny, with the chromids of each genus forming distinct, monophyletic groups (Figure 3). Clear stretches of synteny can be observed when comparing chromids of the genus *Ralstonia* as well as when comparing chromids of the genus *Cupriavidus* (Figure S6). Moreover, the chromids of the genus *Cupriavidus* contain a core gene set of 147 genes, while the core gene set of the *Ralstonia* chromids contain 304 genes (Table 2, Data Set S5). Of the core genes, 58 were common between the two genera (Table 2, Data Set S5). Together, these results are consistent with all chromids of the sister genera *Ralstonia* and *Cupriavidus* being derived from a single replicon acquired by an ancestor at the base of this taxon.

Being present in all examined genomes, the 58 core genes are likely to have been present on the ancestral replicon of the *Ralstonia* and *Cupriavidus* chromids. These 58 core genes include genes involved in oxidative phosphorylation and aerobic respiration, in the pentose phosphate and the Entner-Duodoroff glycolytic pathways (a complete Entner-Duodoroff pathway does not appear to be encoded by the chromosomes), and in nitrite conversion to ammonia, as well as some transcriptional factors, genes involved in resistance, and a cold shock-like protein. Nearly a third of the 58 conserved genes encode products involved in motility (e.g., chemotaxis proteins and flagellum biosynthesis). Although likely not absolutely essential, the genes encoding proteins involved in central carbon metabolism (the Entner-Duodoroff pathway) and chemotaxis are likely to have been relevant in the colonization of new environments; these genes would have provided the possibility to detect and move in response to a chemical stimulus, and they would have increased the metabolic potential of the organism. Hence, these conserved genes may have provided a selective advantage favouring the maintenance of the ancestral plasmid of the modern-day chromids.

In the ParB phylogeny (Figure 3), the chromids of the genera *Ralstonia* and *Cupriavidus* for a distinct group from the chromids of the genera *Burkholderia* and *Paraburkholderia*. Additionally, the *Ralstonia* and *Cupriavidus* chromids are not present in the Rep phylogeny (Figure S4). The absence of these chromids in the Rep phylogeny reflects an inability to detect their replication protein using our HMM based approach and the Rep_3 HMM. This, in turn, suggests that the replication protein of these replicons belongs to a different family than those of the *Burkholderia* and *Paraburkholderia* chromids. Overall, both the ParB and Rep observations support that the ancestral replicon of the *Ralstonia* and *Cupriavidus* chromids was acquired independently of the ancestral replicon of the *Burkholderia* and *Paraburkholderia* chromids.

### *The* Burkholderia cepacia *complex megaplasmids share common ancestry*

In both the ParB and Rep phylogenies, all megaplasmids of the Bcc formed a closely related monophyletic group (Figures 3 and S4). Additionally, synteny analysis revealed long stretches of synteny between the megaplasmids of diverse Bcc species (Figure S7). However, only 18 genes belonged to the core gene set of the megaplasmids (Table 2, Data Set S5). We wondered if the small core gene set may be related to the homologous translocation event that transferred genes between the chromosome and megaplasmid of *Burkholderia cenocepacia* AU-1054 (Guo et al. 2010); however, removing this strain only increased the core genome size to 30 genes. Despite the small megaplasmid core gene set, the results are overall consistent with all megaplasmids of the Bcc being derived from a common ancestral replicon. The high synteny observed in all pairwise comparisons may further suggest that the ancestral replicon was large in size and contained a significant portion of the genes still found on the modern day descendants. Yet, many of these genes may be dispensable, as the small core gene set suggests that different sets of genes have been lost during evolution of each lineage within the Bcc.

Somewhat surprisingly, the megaplasmids of the Bcc were situated close to the chromids of the genera *Burkholderia* and *Paraburkholderia* in both the ParB and Rep phylogenies (Figures 3 and S4). A BLASTp search of the NCBI non-redundant protein database with the ParB protein of the *B. lata* 383 chromid identified the ParB protein of the *B. lata* 383 megaplasmid (74% coverage, and 32% identity). Strikingly, the BLASTp search indicated that the ParB protein of the *B. lata* 383 chromid is more similar to the ParB protein of the Bcc megaplasmids and other replicons of the family *Burkholderiaceae* than it is to the ParB protein of other species. Given this result, it is likely that the megaplasmid of the Bcc and the chromid of the *Burkholderia* are derived from the same ancestral plasmid pool.

### *The* Paraburkholderia *megaplasmids do not share common ancestry*

In contrast to the Bcc, the megaplasmids of the genus *Paraburkholderia* do not form a monophyletic group in either the ParB or Rep phylogenies (Figures 3 and S4). A lack of synteny is observed when comparing megaplasmids belonging to distinct lineages based on the ParB phylogenies (Figure S7), and no set of core genes could be identified (Table 2). It is therefore likely that the megaplasmids of the genus *Paraburkholderia* did not originate from a single ancestral replicon.

### *The ParB protein of the* Burkholderia *chromids have experienced positive selection*

Consistent with the topology of the species phylogeny of Figure 2, the chromids of the Bcc formed a distinct group from the rest of the *Burkholderia* chromids in the ParB phylogeny (Figure 3). A McDonald-Kreitman test (McDonald and Kreitman 1991; Egea et al. 2008) was performed with the ParB proteins of these two groups to evaluate if ParB was experiencing positive selection (i.e., selection for genetic variants). The results are consistent with strong positive selection acting on ParB (Neutrality Index of 0.039, p-value < 0.001; Table S1). We hypothesize that the acquisition of the ancestral megaplasmid at the base of the Bcc clade led to selective pressure for divergence of the ParB proteins of the megaplasmid and resident chromid. A divergence in these proteins would help reduce cross-talk between the partitioning systems of these replicons and promote stable inheritance of both replicons. Indeed, previous studies have demonstrated that the replicon partitioning systems of each replicon in *Burkholderia cenocepacia* (Dubarry et al. 2006; Passot et al. 2012; Du et al. 2016) and *Rhizobium leguminosarum* (Żebracki et al. 2015) specifically recognize and act on their cognate replicon. Presumably, this specialization contributes to the orderly segregation of each replicon that has been observed in *B. cenocepacia* (Du et al. 2016) as well as in *Sinorhizobium meliloti* (Frage et al. 2016).

### Relationship between genome architecture and genome size

It was previously noted that the average size of multipartite genomes is larger than the average size of non-multipartite genomes (Harrison et al. 2010; diCenzo and Finan 2017). Indeed, the multipartite genomes of *Burkholderia, Paraburkholderia*, and *Cupriavidus* are larger than those of *Pandoraea* [whose lifestyle (Data Set S4) is similar to that of the *Burkholderia* (Coenye et al. 2000; Daneshvar et al. 2001), but which lack large secondary replicons] and the obligate endosymbionts of the genus *Polynucleobacter* (Table 1). On the other hand, the multipartite genomes of the genus *Ralstonia* are similar in size to those of the genus *Pandoraea*. In contrast, the *Pandoraea* chromosomes are significantly larger (> 1 Mb) than the chromosomes of all four genera with multipartite genomes (Table 1). Similarly, the chromosomes of the *Burkholderia* strains with a megaplasmid are on average smaller than those without a megaplasmid. The same observation is true for the *Paraburkholderia*. There was little evidence for an effect of megaplasmid presence on chromid size, although we cannot rule out this possibility. Overall, these correlations are consistent with secondary replicons facilitating an enlargement of the genome. However, they also appear to contradict the hypothesis that secondary replicons are required for an enlargement of the genome once the chromosome is at maximum capacity; otherwise, we would expect the chromosomes of the *Pandoraea* to be similar in size to those of the *Burkholderia* and *Paraburkholderia*. Chromids may therefore accumulate numerous genes that otherwise may have been acquired by the chromosome if the chromid was not present.

### Functional characteristics of the *Burkholderiaceae* genomes

Replicon-specific pangenome analyses and Cluster of Orthologous Genes (COG) enrichment analyses (Tatusov et al. 2003) were performed to evaluate the relationships between genome architecture, bacterial lifestyle, and gene functional biases.

#### Increased genetic variability only partially explains inter-replicon functional biases

Several studies have reported functional biases between replicons within a genome, with functions such as transport, metabolism, regulatory functions, and motility commonly enriched on chromids and megaplasmids (see for example, (Heidelberg et al. 2000; Galibert et al. 2001; Paulsen et al. 2002; Chain et al. 2006; diCenzo and Finan 2017)). Secondary replicons are also more genetically variable than the chromosome (see for example, (Holden et al. 2004; Choudhary et al. 2007; Cooper et al. 2010; Epstein et al. 2012; Van Houdt et al. 2012; diCenzo and Finan 2017)). To address if the inter-replicon functional biases are purely a consequence of these replicons being enriched in the accessory pangenome fraction, replicon-specific pangenome functional annotations were performed. Replicon-specific pangenomes were determined with Roary (Page et al. 2015), the representative proteins of each orthologous cluster were annotated with COG terms, and the percentage of accessory proteins (defined as those found in less than 75% of the strains) annotated with each term were compared. The same results were obtained if a threshold of 25% of strains was used (not shown).

As shown in Figure 4, a clear correlation is observed between the abundance of COG categories in the chromosome and chromid or megaplasmid accessory pangenome for all analyzed groups. This suggests a general relationship between the functional biases of each replicon in a genome, consistent with the functional biases of secondary replicons partly being a result of their enrichment in the accessory genome. Nevertheless, inter-replicon functional biases remained at the pangenome level (Figure 4, Table S2). In particular, energy production (category C), transport and metabolism of amino acids, lipids, carbohydrates, and inorganic ions (categories E, G, I, P), secondary metabolism (category Q), transcription (category K), and signal transduction (category T) were enriched in the chromid and/or megaplasmid accessory pangenomes (Table 3, Table S2). Thus, the increased genetic variability of secondary replicons is insufficient to fully explain the inter-replicon functional biases within bacterial multipartite genomes.

**Figure 4.**
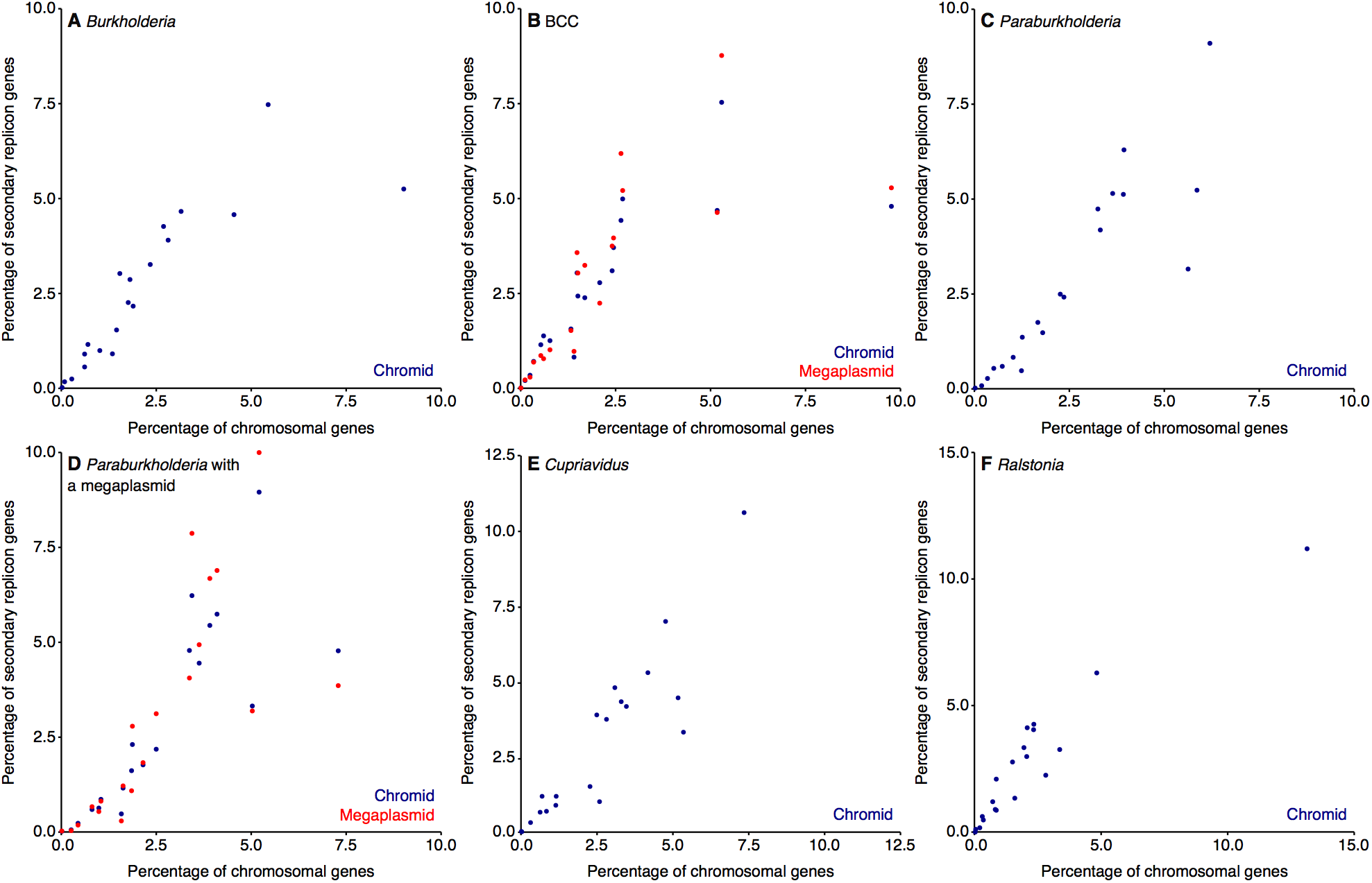
Relationship between COG category abundances in replicon-specific accessory pangenomes. Scatterplots displaying the percentage of proteins in the accessory pangenome of chromids and megaplasmids compared to that of chromosomes are shown. In each plot, the X-axes show the percentage of proteins in the chromosomal accessory genome annotated with a specific COG, while the Y-axes show the percentage of proteins in the secondary replicon (chromid or megaplasmid) accessory genome annotated with a specific COG category. Each point in the plots represent a single COG category. The data underlying these images are provided in Data Set S4. The figures are shown for the following groups of strains: **(A)** all *Burkholderia* strains, **(B)** the *Burkholderia cepacia* complex, **(C)** all *Paraburkholderia* strains excluding strain HKI 454, **(D)** the *Paraburkholderia* strains with a megaplasmid excluding strain HKI454, **(E)** all *Cupriavidus* strains, and **(F)** all *Ralstonia* strains.

**Table 3.** Inter-replicon accessory genome COG functional biases.

| COG<br>Category | *Burkholderia* |  |  | *Paraburkholderia* |  |  | *Cupriavidus* |  |  | *Ralstonia* |  |  |
| --- | --- | --- | --- | --- | --- | --- | --- | --- | --- | --- | --- | --- |
|  | Chromosome | Chromid | Ratio* | Chromosome | Chromid | Ratio* | Chromosome | Chromid | Ratio* | Chromosome | Chromid | Ratio* |
| Total Orthologous Groups † |  |  |  |  |  |  |  |  |  |  |  |  |
| C | 2.69 | 4.26 | 1.58 | 3.94 | 6.29 | 1.60 | 4.76 | 7.00 | 1.47 | 2.08 | 4.12 | 1.98 |
| E | 3.15 | 4.66 | 1.48 | 3.92 | 5.12 | 1.31 | 4.18 | 5.29 | 1.27 | 2.35 | 4.26 | 1.82 |
| G | 2.81 | 3.90 | 1.39 | 3.64 | 5.15 | 1.41 | 2.82 | 3.75 | 1.33 | 2.06 | 2.98 | 1.45 |
| I | 1.76 | 2.26 | 1.28 | 2.36 | 2.42 | 1.02 | 3.09 | 4.80 | 1.55 | 0.86 | 2.09 | 2.43 |
| K | 5.44 | 7.47 | 1.37 | 6.20 | 9.10 | 1.47 | 7.35 | 10.61 | 1.44 | 4.83 | 6.29 | 1.30 |
| P | 2.34 | 3.26 | 1.39 | 3.32 | 4.18 | 1.26 | 3.47 | 4.17 | 1.20 | 2.34 | 4.04 | 1.73 |
| Q | 1.81 | 2.86 | 1.59 | 2.26 | 2.49 | 1.10 | 2.50 | 3.89 | 1.56 | 1.50 | 2.77 | 1.84 |
| T | 1.54 | 3.02 | 1.96 | 3.25 | 4.74 | 1.46 | 3.30 | 4.33 | 1.31 | 1.95 | 3.34 | 1.71 |
| Unique Orthologous Groups ¥ |  |  |  |  |  |  |  |  |  |  |  |  |
| C | 1.34 | 1.92 | 1.43 | 2.13 | 2.61 | 1.22 | 2.63 | 3.14 | 1.20 | 1.62 | 2.98 | 1.84 |
| E | 1.67 | 2.28 | 1.36 | 2.21 | 2.66 | 1.20 | 2.36 | 2.94 | 1.25 | 1.90 | 3.18 | 1.67 |
| G | 1.25 | 1.67 | 1.34 | 1.94 | 2.16 | 1.12 | 1.47 | 1.72 | 1.17 | 1.32 | 2.06 | 1.56 |
| I | 0.87 | 1.04 | 1.19 | 1.18 | 1.16 | 0.99 | 1.62 | 1.96 | 1.21 | 0.71 | 1.54 | 2.16 |
| K | 2.50 | 2.96 | 1.18 | 3.12 | 3.49 | 1.12 | 3.74 | 4.57 | 1.22 | 3.25 | 4.34 | 1.34 |
| P | 1.19 | 1.47 | 1.24 | 1.68 | 1.97 | 1.18 | 1.94 | 2.13 | 1.10 | 1.43 | 2.60 | 1.81 |
| Q | 0.82 | 1.08 | 1.32 | 1.11 | 1.10 | 0.99 | 1.18 | 1.55 | 1.32 | 0.98 | 1.86 | 1.89 |
| T | 0.73 | 1.02 | 1.39 | 1.30 | 1.55 | 1.19 | 1.40 | 1.61 | 1.16 | 1.09 | 1.74 | 1.59 |

\* The ratio is calculated as chromid / chromosome.
† Values represent the percentage of all genes in the replicon-specific pangenome accessory fraction annotated with the indicated COG category (Table S3).
¥ Values represent the number of unique orthologous groups within the indicated COG category in replicon-specific accessory fraction, expressed as a percentage of the total number of genes in the replicon-specific pangenome accessory fraction.

For each of the COG categories mentioned above that are biased between chromosomes and chromids (C, E, G, I, P, Q, K, T), we approximated the functional diversity by counting the number of unique eggNOG orthologous groups (OGs) annotated within each category. Two consistent patterns emerged. First, when considering the number of OGs, the enrichments observed in chromid versus chromosome accessory genomes tended to decrease, but still remained (Table 3). This suggests that the enrichment of these functions in the chromid accessory genome is a combination of the independent acquisition (or duplication) of functionally redundant genes, as well as an extension of the functional repertoire of the replicon. Second, a large fraction (66-88%) of the OGs for each genus were specific to either the chromosomal or chromid accessory genome. This observation further supports the notion that the chromids and chromosomes are functionally divergent, and it suggests that their gene content is derived from a distinct gene pools.

Overall, the above results are consistent with different selective pressures acting on each replicon, thereby selecting for differential functional enrichments on each replicon. This is consistent with results suggestive of differential evolutionary trajectories of each replicon in a genome (Galardini et al. 2013), and the observation of different mutation rates between replicons that is at least partly independent of functional biases (Cooper et al. 2010; Dillon et al. 2015).

#### Functional biases follow lifestyle not taxonomy

The functional contents of the four genera with multipartite genomes were compared to identify the effect of lifestyle. The results revealed that the chromosomal pangenome functional content of the genus *Burkholderia* is similar to that of the genus *Ralstonia*, while that of the genera *Cupriavidus* and *Paraburkholderia* were similar and distinct from the functional content of the other two genera (Figure 5A). A similar pattern was detected for the pangenome functional content of the secondary replicons (Figure 5B). Notably, species of the genera *Burkholderia* and *Ralstonia* are pathogenic, whereas species of the genera *Cupriavidus* and *Paraburkholderia* are environmental isolates (Data Set S4). Hence, the functional content of the species of the family *Burkholderiaceae* appears to be related more to lifestyle than to taxonomy.

**Figure 5.**
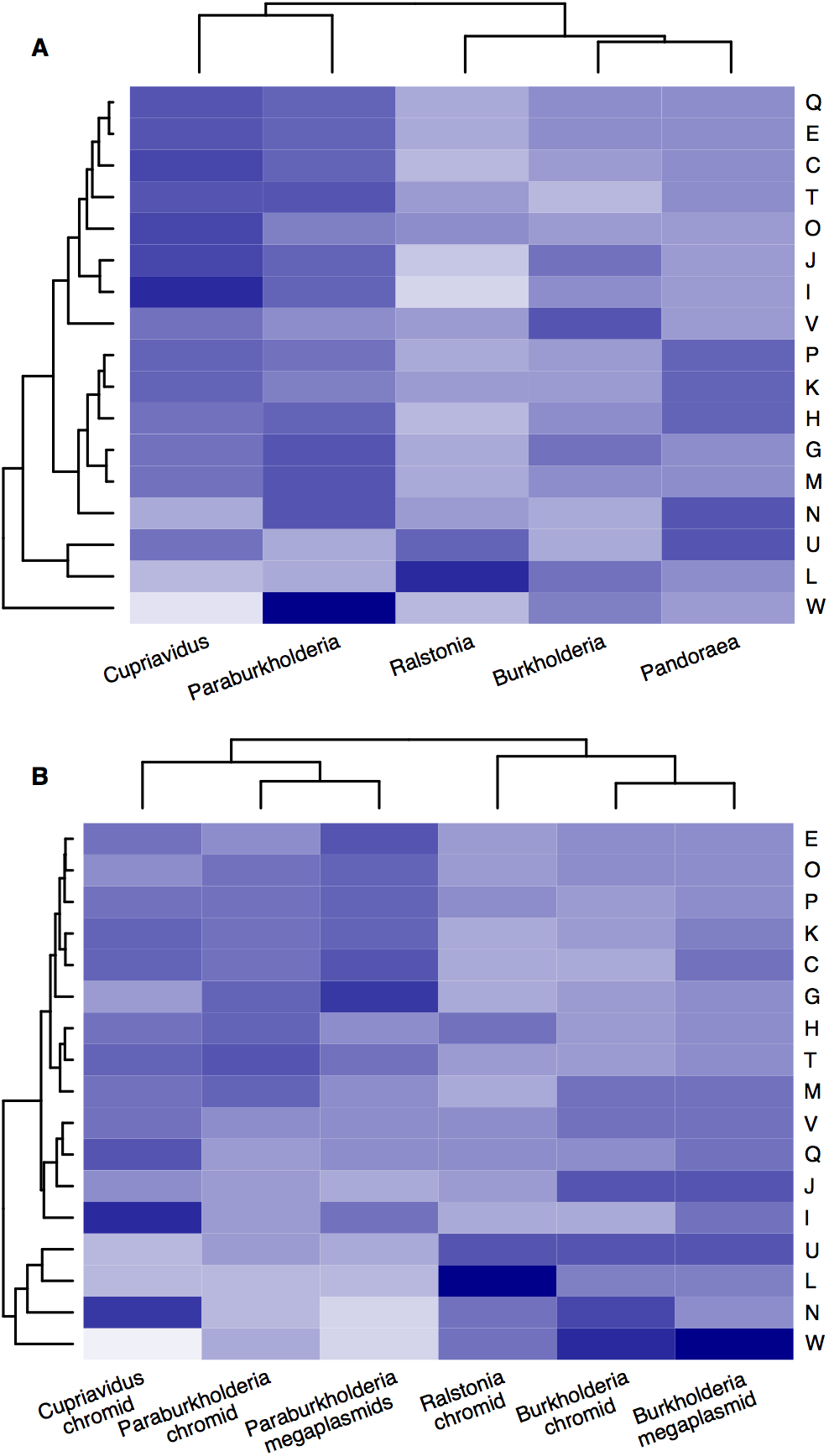
Accessory pangenome COG category abundances. Heatmaps showing the relative abundance of proteins annotated with each COG category in the (**A**) chromosome pangenomes and (**B**) secondary replicon pangenomes is shown. Values shown are the percentage of proteins annotated with the COG category, divided by the average value for all samples shown in the heatmap. Only those COG categories with an unequal distribution between groups is shown, as determined with a Chi-squared test with 10 degrees of freedom and an unadjusted p-value of 0.001. Additionally, COG category S (general functional prediction only) was excluded. Both axes were clustered with hierarchical clustering using average linkage and a Pearson correlation distance.

The observation that the chromosomes of the family *Burkholderiaceae* are functionally biased according to lifestyle suggests that having a chromid is not necessary, *per se*, for adaptation to the distinct niches of these organisms. This is further supported by the pangenome functional content of the chromosomes of the genus *Pandoraea*. This genus lacks secondary replicons but has a lifestyle similar to the genus *Burkholderia*, including being an emerging pathogen found in the lungs of cystic fibrosis patients (Coenye et al. 2000; Daneshvar et al. 2001). It is therefore not surprising that in terms of COG functional abundances, the chromosomal accessory pangenome of the *Pandoraea* was similar to that of *Burkholderia* and *Ralstonia* (Figure 5A and Table S2). However, there did appear to be moderate biases in the chromosomal COG abundances of *Pandoraea* relative to *Burkholderia* and *Ralstonia*, with these biases often being in the direction of those seen for the chromids versus chromosomes. For example, moderate enrichments were observed in COG categories C, K, P, and T (Figure 5 and Table S2). We therefore conclude that a secondary replicon is not an absolute necessity for niche adaptation by these organisms.

#### Chromids appear enriched in environmental adaptation functions

Given that functional biases were detected between pathogenic and environmental genera, and that functional biases were detected between chromosomes and chromids, we examined whether there were similarities between these two sets of biases. As shown in Figure 6A, there is a correlation between the ratio of COG abundance in the chromosomal accessory genome of the *Paraburkholderia* versus the *Burkholderia*, and the ratio of COG abundance in the chromid and chromosomal accessory genomes of these genera. A similar pattern was observed for the genera *Cupriavidus* and *Ralstonia* (Figure 6B). Notably, for both pairs of genera, the observed relationship was stronger for the pathogenic organisms (*Burkholderia* and *Ralstonia*). Thus, at least in the family *Burkholderiaceae*, functions enriched in environmental isolates relative to pathogenic organisms are the same functions enriched on chromids relative to chromosomes. This is especially true for the pathogenic genera, likely as the chromosomes of the environmental strains also contain higher amounts of these genes. These results are therefore consistent with chromids being enriched in environmental, or niche, adaptation genes. However, this result further supports that chromids are not absolutely necessary for the accumulation of niche adaptation genes, and that such genes are likely accumulated on the chromosome even in the presence of a chromid.

**Figure 6.**
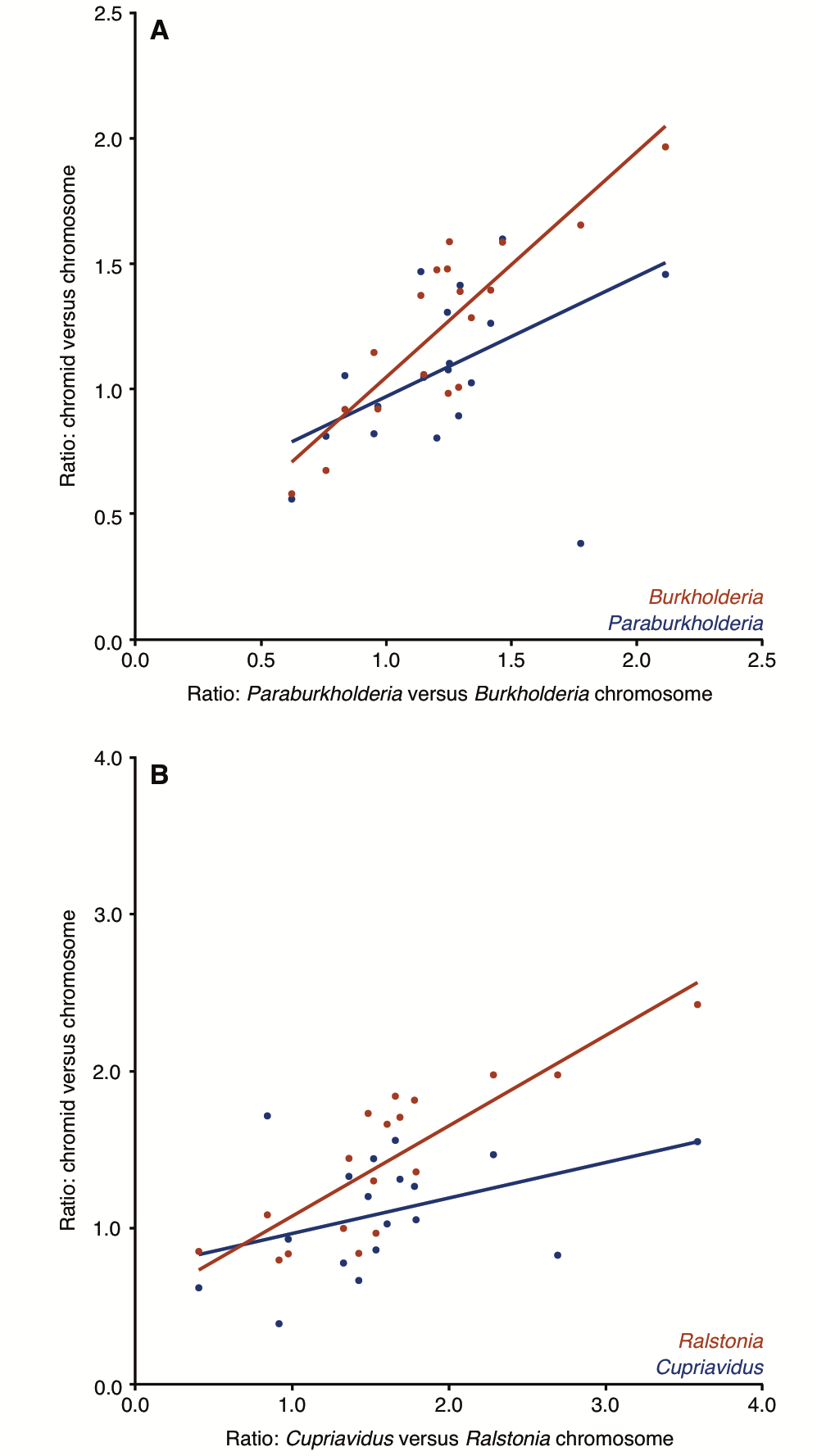
Correlation between replicon and life-style functional biases. Plots display the chromosomal COG functional category abundance ratio for environmental genera versus pathogenic genera (X-axes), compared to the COG functional category abundance ratio for chromids versus chromosomes (y-axes). The axes display the ratio of the percentage of proteins annotated with a COG category. Each point on the plots represents on COG category. COG abundance data is from the replicon-specific accessory genomes. Data points are limited to COG categories with an abundance of at least 0.5%. Lines represent the robust linear regression determined with the ‘rlm’ command of the MASS package for R (Venables and Ripley 2002). (**A**) Data for *Burkholderia* and *Paraburkholderia*. (**B**) Data for *Ralstonia* and *Cupriavidus*.

### Chromids may facilitate increased rates of horizontal gene transfer

It has been suggested that chromid evolution involves significant gene accumulation and replicon enlargement (diCenzo et al. 2014; diCenzo and Finan 2017), and that chromids may permit species to continue to enlarge their genome when the chromosome reaches a maximal size (Slater et al. 2009). We therefore evaluated the evolution of the genome content within the family *Burkholderiaceae*. As described in the Methods, all proteins of the 293 *Burkholderiaceae* strains were grouped into clusters using cd-hit (Li and Godzik 2006), and the presence of each cluster in each ancestral strain was predicted using ancestral state reconstruction, as implemented in the APE package of R (Paradis et al. 2004). We considered two groups of strains: i) the genera *Ralstonia* and *Cupriavidus*, and ii) the genera *Burkholderia* and *Paraburkholderia*. Within each group, the analysis was limited to those clusters found only in the chromosome pangenome or only in the chromid pangenome.

The same overall trend was observed in both groups of strains (Figure 7). Moving from the base of the tree to the extant strains, there was generally an increase in the number of genes found in the ancestral strains that are also present in the modern-day strains. This was true for both the chromosomal genes and for the chromid genes. However, the proportional increase in the number of chromid genes outpaced that in the rate of gain of chromosomal genes (Figure S8), an observation that was particularly pronounced in the *Burkholderia* and *Paraburkholderia*. These observations are consistent with the ancestral chromid replicons carrying a relatively small portion of the genes currently found on the modern-day descendants. Additionally, these results are consistent with secondary replicons more rapidly acquiring (and possibly losing) genes through horizontal transfer than chromosomes, contributing to increased genome plasticity.

**Figure 7.**
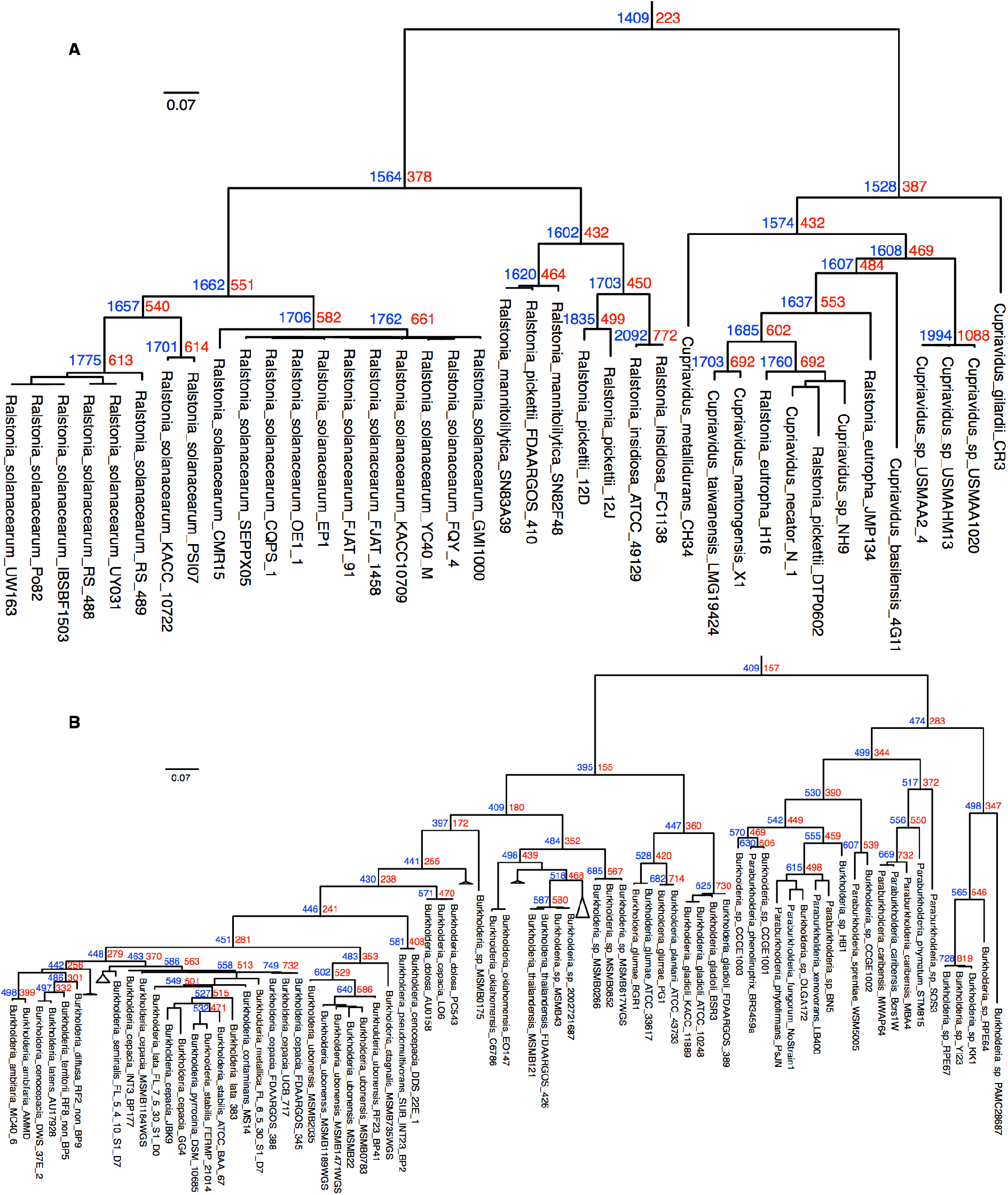
Ancestral state reconstruction of genome content. Sub-trees of the phylogeny shown in Figure 2 are shown for **(A)** the genera *Cupriavidus* and *Ralstonia*, and **(B)** the genera *Burkholderia* and *Ralstonia*. At each node (space permitting), the number of genes found to the chromosomal pangenome (blue) and the chromid pangenome (red) of the extant strains of the tree that are predicted to have been present in that ancestor are indicated.

## CONCLUSIONS

It has been noted elsewhere that ~ 10% of sequenced bacterial species contain a chromid (Harrison et al. 2010; diCenzo and Finan 2017), suggesting that the presence of a chromid is uncommon but not rare. In our analysis of the β-proteobacteria, we examined 213 named species, of which 49 (23%) contained a chromid (Figure 1). Notably, all but one of these were within the family *Burkholderiaceae*, and the data suggest that the chromids in this family arose from just two ancestral plasmids; however, it remains possible that the transition from plasmid to chromid may have occurred multiple times. It has also been proposed that the chromids of the *Rhizobiaceae* family, which account for the majority of the chromids in the a-proteobacteria, arose from a single ancestral plasmid (Slater et al. 2009). Thus, we conclude that it is likely that the majority of chromids in sequenced strains arose from a few parental replicons, and that the emergence of chromids is likely rarer than suggested by the 10% prevalence.

On average, bacteria containing a chromid have a larger total genome than bacteria lacking chromids, and the difference in genome size can be attributed to the presence of the chromid (Harrison et al. 2010; diCenzo and Finan 2017). This observation has led to the suggestion that the role of chromids is to allow further genome enlargement than is possible with only a chromosome; however, some evidence is also inconsistent with this hypothesis (Egan et al. 2005; diCenzo and Finan 2017). Here, we observed that genera with a chromid had a larger genome than related genera without a chromid (Table 1). However, the chromosomes of these genera tended to be smaller than the chromosomes of a related genus (*Pandoraea*) without a chromid (Table 1). Similarly, α-rhizobia with a chromid contain a smaller chromosome than α-rhizobia lacking chromids (MacLean et al. 2007). This suggests that, at least in some cases, species with a chromid do not contain a chromosome that has reached a maximal size limit. We therefore conclude that the presence of a chromid cannot be solely explained by a need for a mechanism to increase genome content, although chromids may facilitate this process.

Another theory is that chromids (and megaplasmids) serve niche specialized roles, and their evolution was driven by adaptation to novel environments (Chain et al. 2006; Galardini et al. 2013; diCenzo et al. 2014). Consistent with this, our results indicate that the chromid pangenomes are enriched in COG functional categories (Table 3) that are often associated with environmental adaptation (Tamames et al. 2016; Cobo-Simón and Tamames 2017). On the other hand, these same functions tended to be enriched on chromosomes of environmental isolates relative to pathogenic isolates (Figure 5). They also appeared somewhat enriched on the chromosomes of the genus *Pandoraea*, which lacks a chromid, relative to the *Burkholderia*, which contain a chromid (Figure 5A). Hence, while secondary replicons may indeed be niche specialized, chromids are not essential for adapting to these environments, and the same functions can instead be accumulated by chromosomes in the absence of secondary replicons.

We therefore propose a modification of the theory that the primary advantage of secondary replicons is related to niche adaptation. We suggest that the main advantage provided by a secondary replicon is increased genetic malleability, which consequently results in environmental specialization. If acquisition of the secondary replicon corresponds with the ability to inhabit a new environment, the cell will experience new selective pressures selecting for the acquisition of a distinct repertoire of genes. If secondary replicons are more genetically malleable than chromosomes, these new genes would be preferentially accumulated by the secondary replicon, thereby resulting in its enlargement and a replicon specialized for adaptation to a particular environment. In support of this, hot spots for horizontal gene transfer have been observed in bacterial genomes (Oliveira et al. 2017), and others have noted that secondary replicons show increased genetic variability (Holden et al. 2004; Choudhary et al. 2007; Cooper et al. 2010; Epstein et al. 2012; Van Houdt et al. 2012; diCenzo and Finan 2017), increased rates of evolution (Chain et al. 2006; Cooper et al. 2010; Galardini et al. 2013; Dillon et al. 2015; Dillon et al. 2017), and distinct evolutionary trajectories (Galardini et al. 2013). Moreover, our data was consistent with secondary replicons accumulating new genes more rapidly than chromosomes (Figures 7, S8). Thus, the main benefit of secondary replicons may be to increase the ability of the cell to expand its genome content through horizontal gene transfer, in turn resulting in a niche specialized replicon, as opposed to the niche adaptation directly being the prime advantage.

A remaining question is why secondary replicons may be more genetically malleable. It was suggested that chromosomal hot spots for horizontal gene transfer are the result of recombination mechanisms and natural selection (Oliveira et al. 2017). The same mechanisms may explain the increased genetic malleability of secondary replicons. The topology of chromosomes and secondary replicons may differ (Val et al. 2016). Additionally, secondary replicons, like chromosomes, encode nucleoid-associated proteins (NAPs) (Shintani et al. 2015), possibly influencing the relative level of DNA compaction of the different replicons. Thus, secondary replicons may be more available for integration of horizontally acquired DNA through recombination. Secondary replicons, especially in their early evolution, contain few housekeeping genes (Harrison et al. 2010); for example, Tn-seq screens with *S. meliloti* and *B. cenocepacia* have identified 10-fold to 50-fold enrichments of essential genes on the chromosome relative to the secondary replicons (Higgins et al. 2017; diCenzo et al. 2018). As such, integration of new DNA within secondary replicons have a lower probability of disrupting an important gene. Moreover, integration of genes into a secondary replicon may result in lower expression than if integrated into the chromosome (Morrow and Cooper 2012), and lowly expressed, horizontally acquired genes are more likely to be maintained in the genome than highly expressed genes (Park and Zhang 2012). Thus, cells acquiring the foreign DNA on a chromosome may be more likely to be lost from the population than those acquiring the DNA on a secondary replicon, due to the fitness costs of disrupting important genes (Elena et al. 1998) and the higher expression (Morrow and Cooper 2012).

## SUPPLEMENTARY MATERIALS

**File S1.** Contains Tables S1 and S3, and Figures S3-S8.

**Table S1.** Output of McDonald-Kreitman test for positive selection on the *Burkholderia* ParB proteins. Based on 146 ParB proteins from *Burkholderia* strains outside of the Bcc (species 1) and 53 ParB proteins from the Bcc (species 2).

**Table S2.** COG functional analysis data.

**Table S3.** COG category descriptions.

**Figure S1. Phylogeny of the β-proteobacteria.** A RAxML maximum likelihood phylogeny of 960 β-proteobacteria based on 24 highly conserved proteins (see methods). A colour legend is provided in the figure. The branch lengths for one clade were reduced for presentation purposes; this clade is in dashed lines, and a separate scale is provided. Bootstrap values based on 1008 replicates are provided where space permits. The original figure, the Newick formatted phylogeny, and the annotation file are provided in Data Set S6.

**Figure S2. ANI matrix of the family *Burkholderiaceae*.** This figure displays the pairwise average nucleotide identity (ANI) values for each pair of genomes as calculated with FastANI. Values were clustered along both axes using hierarchical clustering with average linkage.

**Figure S3. Size distribution of the replicons in the family *Burkholderiaceae*.** A histogram is provided that displays the size distribution of all non-chromosomal replicons (chromids, megaplasmids, plasmids) from the 293 *Burkholderiaceae* strains.

**Figure S4. Phylogenetic relationship of replicons based on a replication protein.** A RAxML maximum likelihood phylogeny of the replicons of the family *Burkholderiaceae* based on the amino acid sequence of the Rep_3 family partitioning protein. The chromids of *Cupriavidus* and *Ralstonia* are not included as their replication protein does not belong to the Rep_3 protein family. Each taxon is named according to the NCBI accession number of the replicon. A colour legend is provided in the figure. Bootstrap values based on 756 replicates are provided where space permits. The original figure, the Newick formatted phylogeny, and the annotation file are provided in Data Set S6.

**Figure S5. Synteny of the chromids of the genera *Burkholderia* and *Paraburkholderia*.** Dot plots were prepared with the MUMMER package to visualize regions of synteny between **(A-C)** three representative chromids of the genus *Burkholderia*, and **(D-F)** three representative chromids of the genus *Paraburkholderia*. Alignments are coloured according to whether they are in the forward (blue) or reverse (red) direction.

**Figure S6. Synteny of the chromids of the genera *Cupriavidus* and *Ralstonia*.** Dot plots were prepared with the MUMMER package to visualize regions of synteny between **(A-C)** three representative chromids of the genus *Cupriavidus*, and **(D-F)** three representative chromids of the genus *Ralstonia*. Alignments are coloured according to whether they are in the forward (blue) or reverse (red) direction.

**Figure S7. Synteny of the megaplasmids of the genera *Burkholderia* and *Paraburkholderia*.** Dot plots were prepared with the MUMMER package to visualize regions of synteny between **(A-C)** three representative megaplasmids of the *Burkholderia cepacia* complex, and **(D-F)** three representative megaplasmids of the genus *Paraburkholderia*. Alignments are coloured according to whether they are in the forward (blue) or reverse (red) direction.

**Figure S8. Relationship between chromosome and chromid size in ancestral *Burkholderiaceae* strains.** Scatter plots display the ratio between gene count of the chromid versus gene count of the chromosome (Y-axes) versus the gene count of the chromosome (X-axes). Data is based on the ancestral state reconstruction data summarized in Figure 7, and the plots display the data for all nodes. **(A)** Data for the genera *Cupriavidus* and *Ralstonia*. **(B)** Data for the genera *Burkholderia* and *Paraburkholderia*.

**Data Set S1. Classification of all replicons of all strains from the β-proteobacteria.** Information of all replicons from all strains within the β-proteobacteria available through NCBI with a genome status of “finished” or “chromosome”. Replicon type was determined based on GC content and dinucleotide composition as described in the Materials and Methods. GC_difference refers to the difference in GC content of the replicon relative to the GC content of the chromosome of the same strain. Dinucleotide_distance refers to the dinucleotide relative abundance distance between the replicon and the chromosome of the same strain. Representative_genome indicates if the strain is (1) or is not (0) included within the list of representative β-proteobacteria.

**Data Set S2. Classification of all replicons of the representative β-proteobacterial strains.** Information of all replicons from a representative set of β-proteobacteria. Replicon type was determined based on GC content and dinucleotide composition as described in the Materials and Methods. GC_difference refers to the difference in GC content of the replicon relative to the GC content of the chromosome of the same strain. Dinucleotide_distance refers to the dinucleotide relative abundance distance between the replicon and the chromosome of the same strain.

**Data Set S3. Classification of all replicons from all strains of the family *Burkholderiaceae*.** Information of all replicons from all strains within the family *Burkholderiaceae* available through NCBI with a genome status of “finished” or “chromosome”. Replicon type was determined based on GC content and dinucleotide composition as described in the Materials and Methods, and replicon assignments were then manually refined based on apparent common ancestry. GC_difference refers to the difference in GC content of the replicon relative to the GC content of the chromosome of the same strain. Dinucleotide_distance refers to the dinucleotide relative abundance distance between the replicon and the chromosome of the same strain.

**Data Set S4. Lifestyles of the species of the family *Burkholderiaceae*.** A table indicating the niches inhabited by each species of the family *Burkholderiaceae*. The table is based on information available in the GOLD database, the Global Catalogue of Microorganisms, and a literature search.

**Data Set S5. Core, conserved genes of the chromids and megaplasmids.** Contains a list of orthologous groups and associated functions for the conserved genes of i) the chromid of the genera *Burkholderia* and *Paraburkholderia*, ii) the chromid of the *Cupriavidus* and *Ralstonia*, and iii) the megaplasmid of the Bcc.

**Data Set S6. Data underlying the phylogenies of this study.** For each of the phylogenies provided in this study, three files are provided within this data set. These include the original, high-quality figure (“image” file), the Newick formatted tree including bootstrap values (“newick” file), and the annotation file used for colour coding the tree (“annotation” file).

